## Supplementary Material for "Sequence-based prediction of the solubility of peptides containing non-natural amino acids"

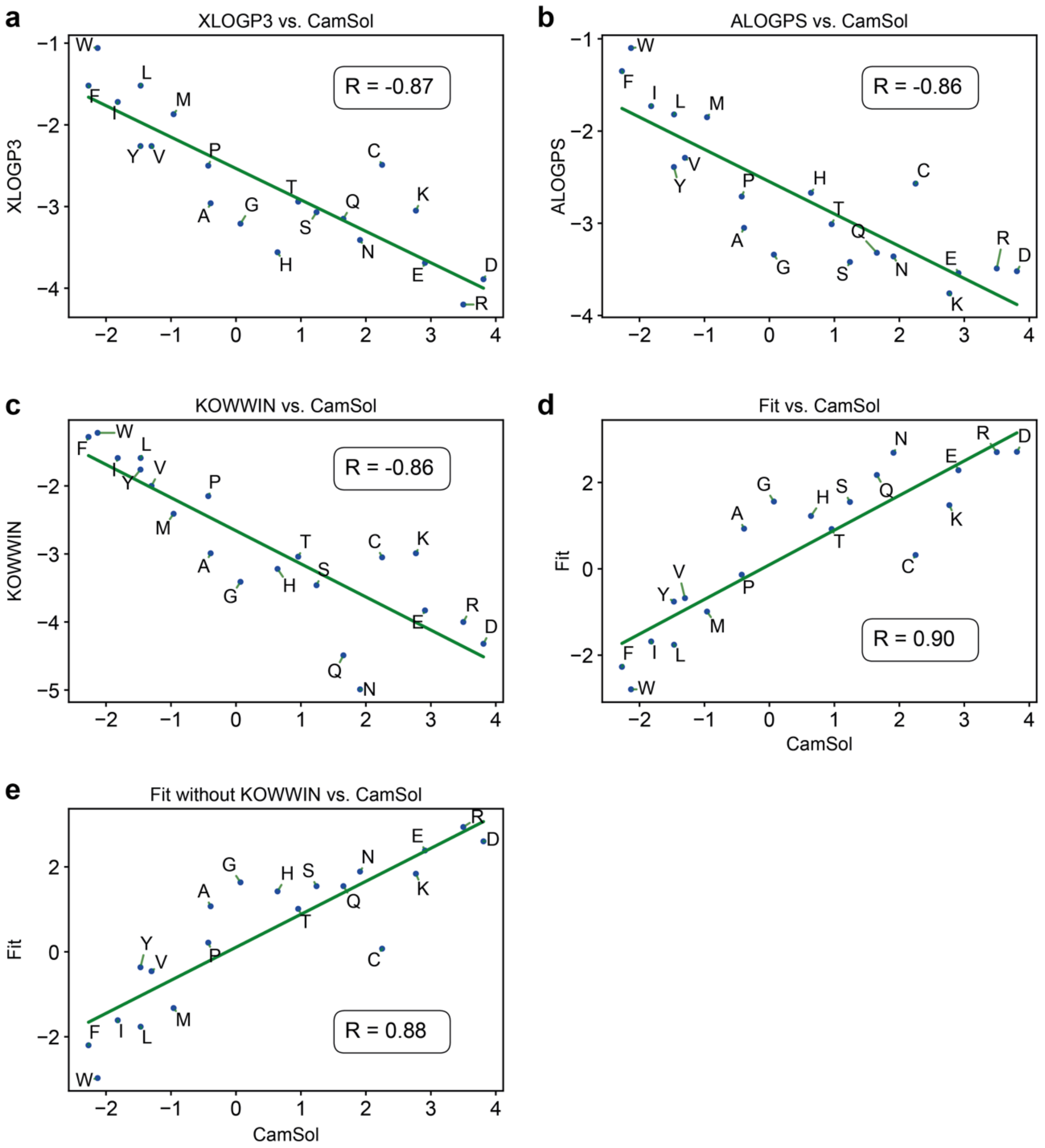


**Figure S1. Comparison of different hydrophobicity predictors with CamSol hydrophilicity values.** Correlation between the hydrophobicity values of XLOGPS **(a)**, ALOGP3 **(b)** and KOWWIN **(c)** and the tabulated hydrophilicity values used in CamSol. **(d)** Values from all three predictors were used to fit a linear regression model to the CamSol hydrophilicity values and then plotted to show the correlation between the fit and the tabulated CamSol hydrophilicity values (Pearson's coefficient of correlation = 0.9). **(e)** Fit without KOWWIN against CamSol hydrophilicity values (Pearson's coefficient of correlation = 0.88).

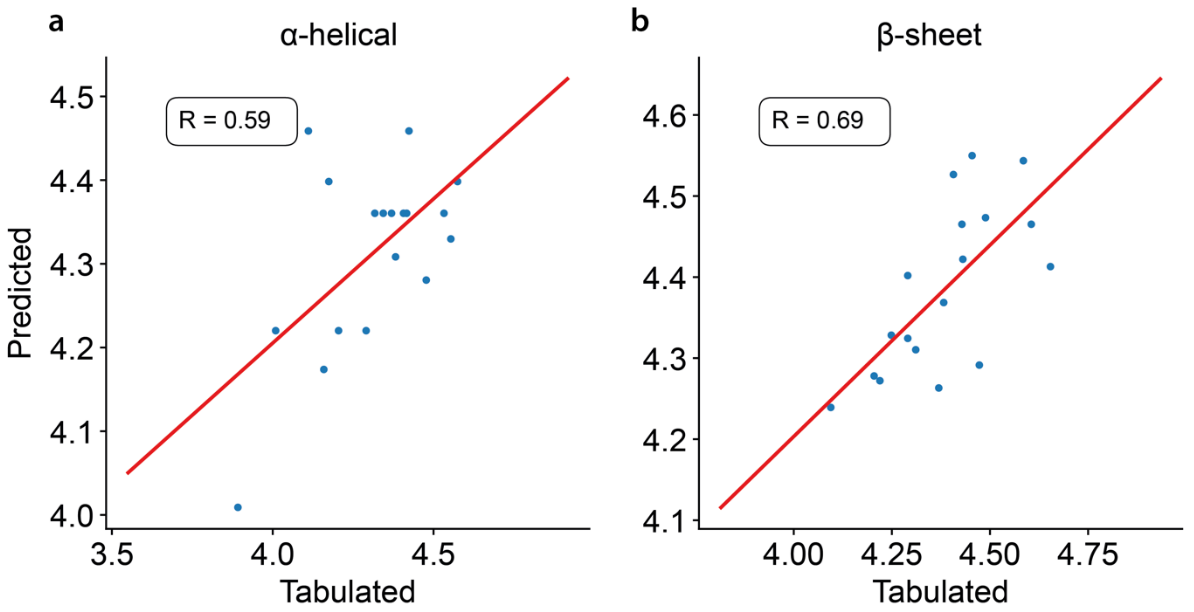


**Figure S2. Correlation between predicted and tabulated secondary structure propensities.** The secondary structure propensity predictor predicts the α-helical propensity **(a)** and β-sheet propensity **(b)** for the 20 natural amino acids with Pearson's coefficients of correlation of 0.59 and 0.69, respectively.

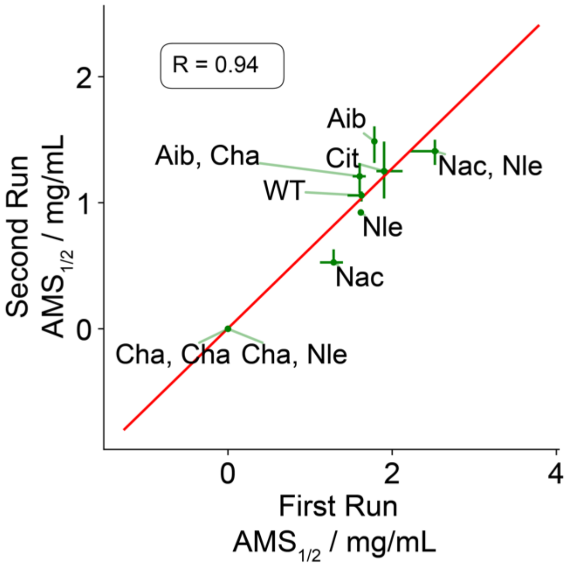


**Figure S3. Comparison of independent AMS runs.** To verify that replacing PEG with AMS continues to yield reliable and replicable results we performed two independent solubility experiments with AMS on two separate days. The correlation indicates that replacing PEG with AMS is feasible.

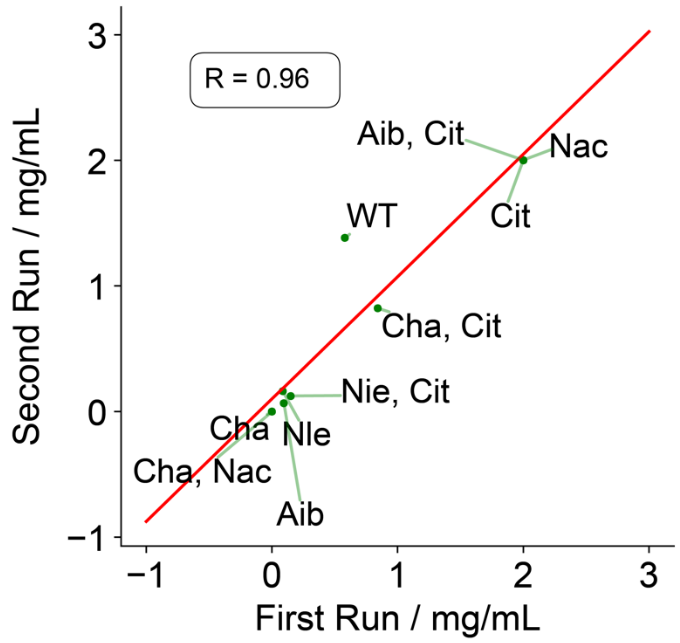


**Figure S4. Comparison of independent ultracentrifugation runs.** To verify the reproducibility of the ultracentrifugation method we performed two independent experiments on different days. The high correlation shows that the results from the ultracentrifugation assay are reliable.

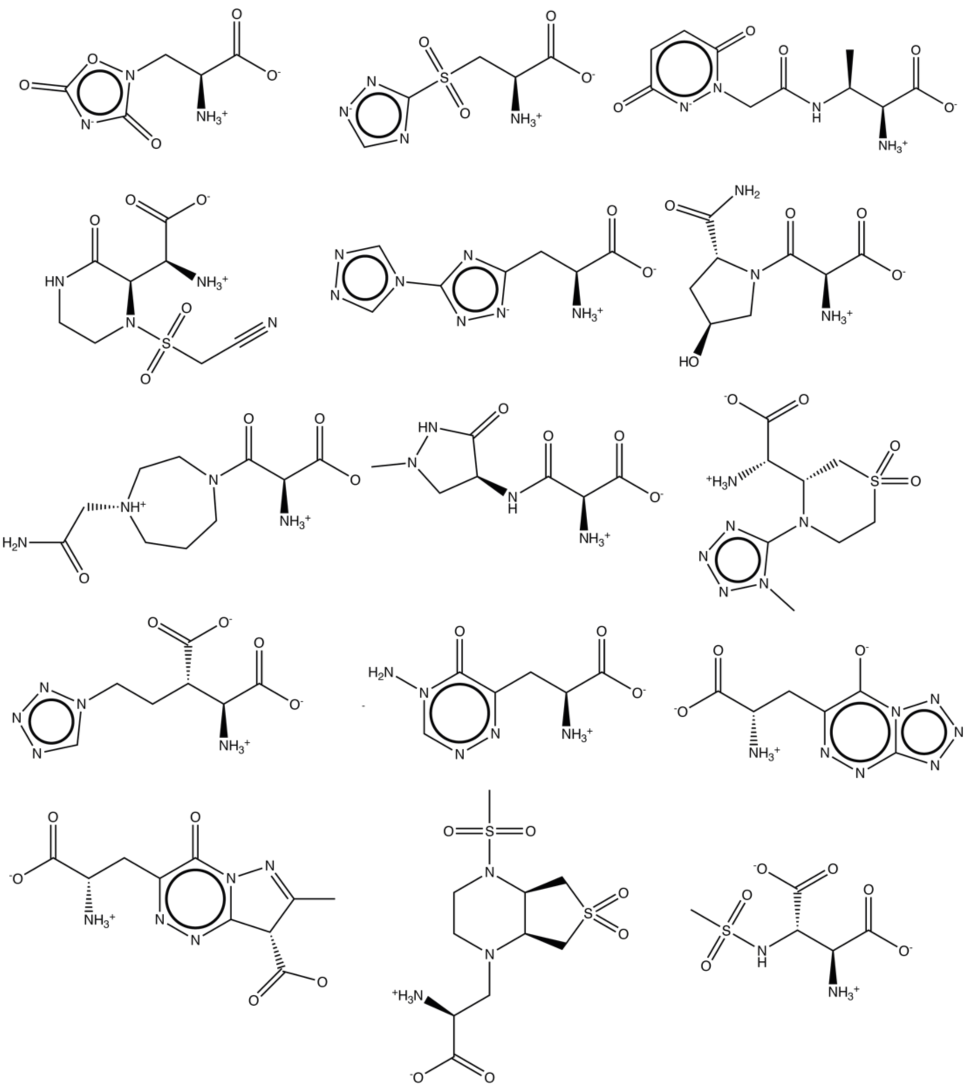

**Figure S5. Selection of mAAs that most effectively promote solubility.** By analysing the positive tail end of the solubility prediction distribution, we found that most mAAs that have a positive effect on solubility contain many hydrogen bonding atoms such as nitrogen and oxygen.

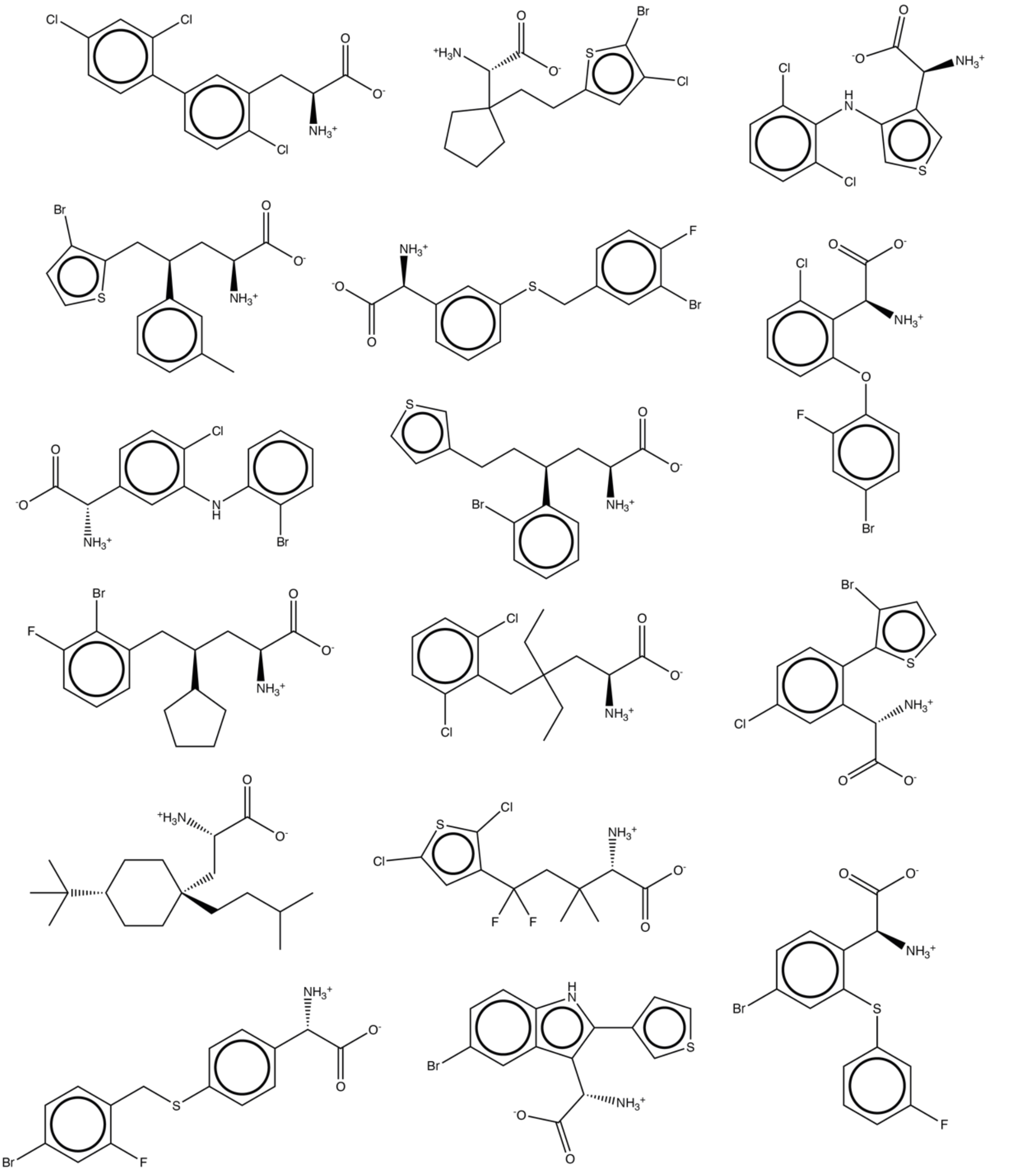

**Figure S6. Selection of mAAs that most effectively decrease solubility.** By analysing the negative tail end of the solubility prediction distribution, we found that most mAAs that have a negative effect on solubility contain several aromatic rings and often halogens such as chlorine or bromine.
